# When Does Mutualism Offer a Competitive Advantage? A Game-Theoretic Analysis of Host-Host Competition in Mutualism

**DOI:** 10.1101/2021.07.13.452216

**Authors:** Abdel H. Halloway, Katy D. Heath, Gordon G. McNickle

**Affiliations:** Department of Plant Biology, University of Illinois at Urbana-Champaign, 505 South Goodwin Ave., Urbana, IL 61801; Department of Botany and Plant Pathology, Purdue University, 915 W. State Street, West Lafayette, IN, 47907, USA; Purdue Center for Plant Biology, Purdue University, West Lafayette, IN, 47907, USA

## Abstract

Plants due to their non-motile nature rely heavily on mutualistic interactions to obtain resources and carry out services. One key mutualism is the plant-microbial mutualism in which a plant trades away carbon to a microbial partner for nutrients like nitrogen and phosphorous. Plants show much variation in the use of this partnership from the individual level to entire lineages depending upon ecological, evolutionary, and environmental context. We sought to determine how this context dependency could result in the promotion, exclusion, or coexistence of the microbial mutualism by seeing if and when the partnership provided a competitive advantage to the plant. To that end, we created a simple 2 × 2 evolutionary game in which plants could either be a mutualist and pair with a microbe or a non-mutualist and forgo the partnership. This model included nutrients freely available to the plant, nutrients obtained only through mutualism with microbes, the cost of producing roots, the cost of trade with microbes, and the size of the local competitive neighborhood. Not surprisingly, we found that mutualism could offer a competitive advantage if its net benefit was positive. Coexistence between strategies is possible though due to competition between mutualists over the microbially obtained nutrient. In addition, the greater the size of the local competitive neighborhood, the greater the region of coexistence but only at the expense of mutualist fixation (non-mutualist fixation was unaffected). Our model, though simple, shows that plants can gain a competitive advantage from using a mutualism depending upon the context and points to basic experiments that can be done to verify the results.

## Introduction

Mutualisms are an important aspect of plant ecology. The non-motile nature of plants means they frequently rely on other organisms to carry out functions such as seed dispersal, pollination, and nutrient acquisition (Howe and Westley 1990). Two key nutrient acquisition strategies for plants are the microbial symbioses with mycorrhizae (in 80% of plant species and 92% of plant families (Simon *et al*. 1993; Wang and Qiu 2006)) and symbiotic nitrogen-fixing bacteria (in a smaller subset of families (de Faria *et al*. 1989; Sprent 2005)). In these mutualisms, the plants trade carbon in the form of carbohydrates and lipids while receiving nutrients like nitrogen and phosphorous (Hawkins *et al*. 2000; Hodge *et al*. 2001; Sessitsch *et al*. 2002; Sawada *et al*. 2003; Leigh *et al*. 2009). Across the plant kingdom, the commonality of partnering with microbial mutualists implies that doing so often offers a fitness benefit to plants (Hartnett *et al*. 1993). However, it is also known that the costs and benefits of mutualism depend upon ecological and evolutionary factors such as nutrient availability and genotype (Peng *et al*. 1993; Heath and Tiffin 2007; Bronstein 2009; Chamberlain *et al*. 2014; Lu and Hedin 2019). These variations in benefits can have knock-on effects at larger scales leading to the variation in the presence or absence of the mutualist partnership among lineages (de Faria *et al*. 1989; Werner *et al*. 2015; Maherali *et al*. 2016). In this paper, we sought to determine how ecological, and environmental context could promote or exclude the microbial mutualistic partnership and ultimately lead to its evolution in a species.

To understand how context determines evolution of microbial mutualisms, we turned to mathematical analysis. Mathematical analysis has been widely used to understand the evolution and persistence of mutualism (Noë and Hammerstein 1995; Ferriere *et al*. 2002; West *et al*. 2002; Hoeksema and Kummel 2003; Akçay and Roughgarden 2007; Akçay and Simms 2011). Typically, the focus of these models has been on the stability and maintenance of interactions between partners, the host and the symbiont, with reasons such as partner selection (West *et al*. 2002; Akçay and Roughgarden 2007; Akçay and Simms 2011) and spatial structure given (see (Wilson *et al*. 2003) for a model of seed dispersal). That said, intraspecific individual competition is a necessary component of evolution by natural selection as the adaptations of more fit individuals become common within the population (Darwin 1859). As such, mutualism must also offer a competitive advantage to a host if it is to evolve (Jones *et al*. 2012). We wanted to explore how host-host competition affects the evolution of mutualism. To do so, we turned to evolutionary game theory. Originally developed to understand animal behavior, evolutionary game theory is a mathematical framework that examines how strategies perform, in terms of fitness, against other interacting strategies (Maynard-Smith and Price 1973; Geritz *et al*. 1998; Brown 2016). It has been applied widely across taxa; for plants, it has been used to understand properties such as defense against herbivory and biomass allocation with competition (Givnish 1982, 1995; Augner *et al*. 1991; McNickle *et al*. 2016). Recently, evolutionary game theoretic host-host competition has been used to understand the global distribution of nutrient acquisition strategies (Lu and Hedin 2019). Viewing the partnership with microbes (and its complement, non-partnership) as strategies in an evolutionary game narrows our focus to just the competitive interactions between hosts and the ecological and environmental contexts that benefit one strategy over the other.

To this end, we created a simple 2 × 2 matrix game to determine how nutrient availability, frequency of alternate strategies, and competitor density may (or may not) offer an intraspecific competitive advantage to a plant that partners with a microbe to obtain nutrients. In our model, we assume that the mutualism partnership is itself a strategy, the equivalent of a functional trait (Violle *et al*. 2007), where a plant can either be a non-mutualist and only acquire benefits from freely available nutrients in the soil or be a mutualist and receive additional benefits from microbially obtained nutrients. All plants must pay a cost to acquire the freely available nutrients with mutualists paying an additional cost for the microbially obtained nutrients. Besides these four parameters, we also included local competitor number as a parameter to see how density-dependence may influence selection (Clarke 1972). We analyzed our game for the fixation of either strategy as well as coexistence of both strategies within a population. We discuss what our results mean for the evolution of and variation in mutualist strategies in plant-microbe systems.

## Model Analysis

### Competition with one plant

In our model, we start out by assuming there are two pools of nutrients available to a plant: one that is freely available *AN* and one that is only obtained through microbial mutualism *MN*. These nutrients provide fitness benefits of *B*_*AN*_ and *B*_*MN*_ respectively to a plant. Some proportion of the population is the genotype of plants with the ability to partner with microbial mutualists while the remainder is made up of the genotype that cannot; we hereafter refer to those genotypes as mutualists and non-mutualists respectively. Non-mutualist plants only get the fitness benefit from the freely available nutrients while mutualists get fitness benefits from both freely available nutrients and microbially obtained nutrients. All plants must produce roots to obtain the freely available nutrient at a cost of *c*_*r*_. Mutualists however have to pay an additional fitness cost *c*_*t*_ to obtain the microbial nutrients due to trade and other mechanisms (e.g., allocation of biomass to nodules in the case of rhizobia mutualism). Finally, we begin our analysis by assuming only two plants compete at a given instant with each plant having equal competitive ability. From these assumptions, we construct the following fitness matrix for each type of plant:

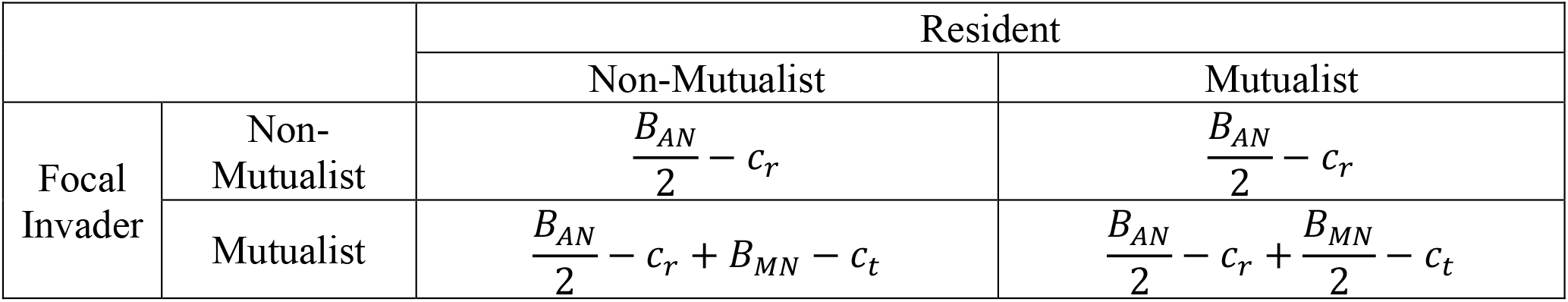

|  |  | Resident |  |
| --- | --- | --- | --- |
|  |  | Non-Mutualist | Mutualist |
| Focal Invader | Non-Mutualist | $\frac{B_{AN}}{2} - c_r$ | $\frac{B_{AN}}{2} - c_r$ |
| | Mutualist | $\frac{B_{AN}}{2} - c_r + B_{MN} - c_t$ | $\frac{B_{AN}}{2} - c_r + \frac{B_{MN}}{2} - c_t$ |

Since all individuals have access to freely available nutrients and must produce roots, all individuals get a net fitness benefit of 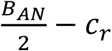 regardless of strategy as competition over the freely available nutrients means each individual only receives half of the potential fitness benefit from that pool of resources. If a mutualist competes with a non-mutualist, the mutualist gets the full benefit of the microbially obtained nutrients while paying the cost of trade *B*_*MN*_ − *c*_*t*_; however, when competing with another mutualist, both compete over and therefore equally share the microbially obtained nutrients leading to a net benefit of 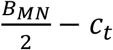.

Since all individuals receive the exact same fitness benefit from the freely available nutrient and pay the exact same cost for the roots 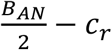, these terms can be removed to arrive at the simpler payoff matrix below:

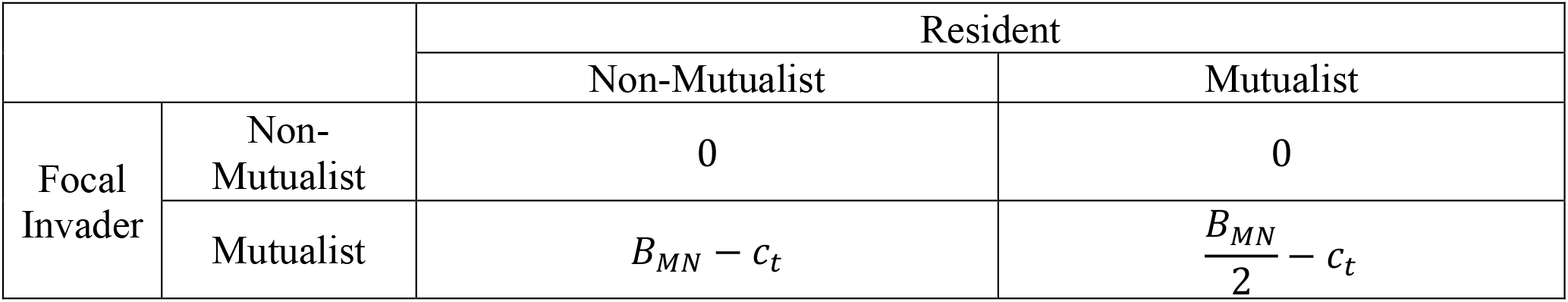

From this simplified matrix, we can quickly arrive at conditions for fixation of mutualist or non-mutualist varieties. Specifically, if the cost of trade outweighs the total benefit of microbially obtained nutrients *c*_*t*_ > *B*_*MN*_, then mutualist do worse, and non-mutualism is the dominant strategy (Fig 1a,b). This is intuitive and true of any trait: when the fitness costs outweigh the benefits, no trait should be favored by natural selection. However, if the benefits of microbially obtained nutrients after competition with other mutualist plants in the population is greater than the cost of trade 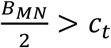, then mutualists always do better and so become the dominant strategy (Fig 1 c,d). Interestingly, the difference between *B*_*MN*_ and 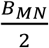 creates a region of the fitness landscape where mutualists and non-mutualists can coexist within a population. Indeed, if the total benefit of microbially obtained nutrients is greater than the cost of trade but the benefit of microbially obtained nutrients under competition is lower than the cost of trade (i.e. 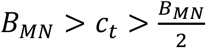), then both genotypes coexist in the same shared space (Figure 1e,f). Solving for the equilibrium proportion of mutualists in the population gives 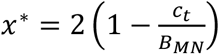 (Figure 2). This coexistence point is a stable equilibrium (Figure 1f).

**Fig. 1.**
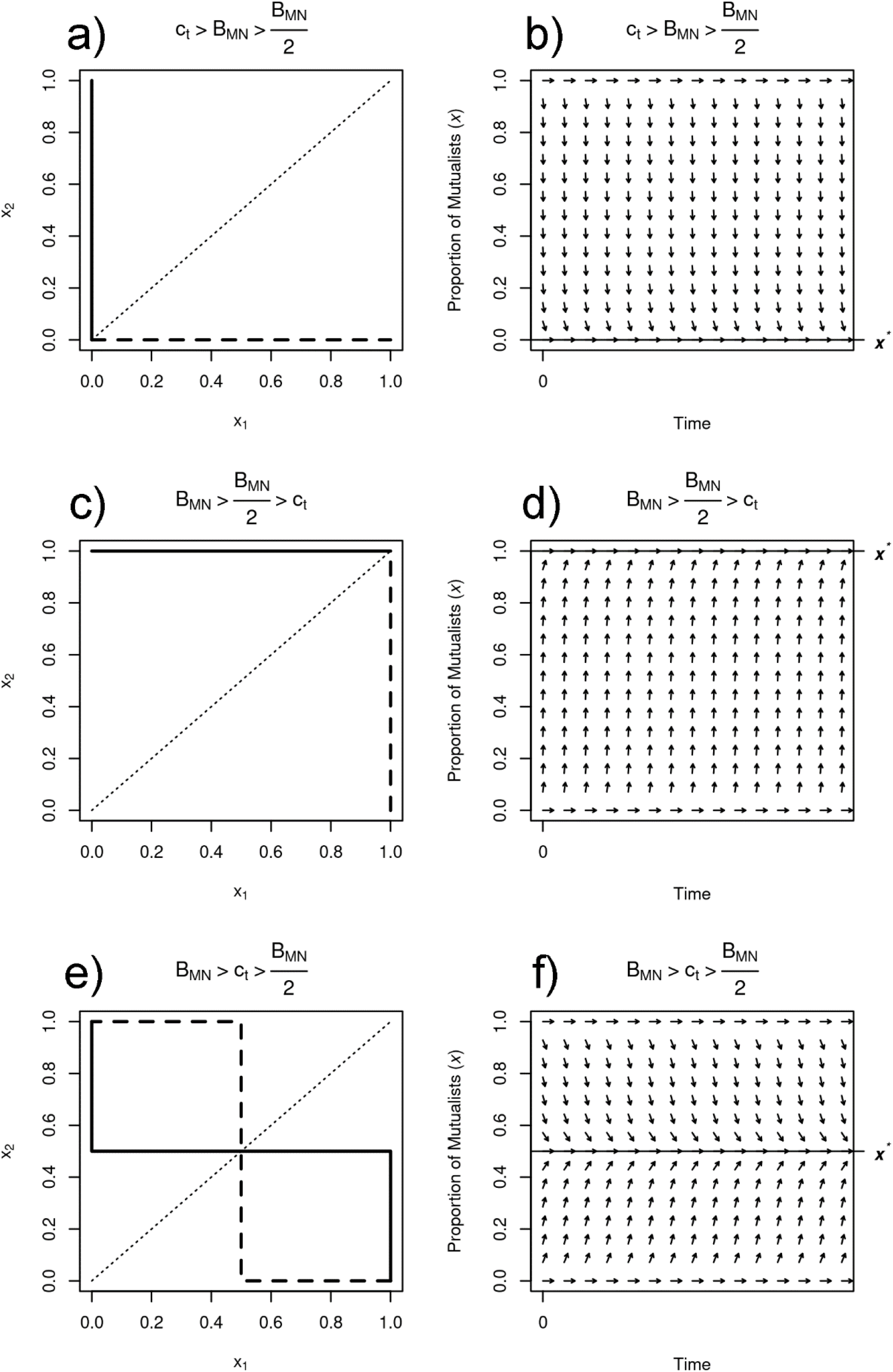
Evolutionary dynamics as seen through best response curves (a, c, e) and directional fields (b, d, f) for the three qualitatively different scenarios. In the first scenario (a, b), the cost of mutualism outweighs any benefit regardless of the opposing player’s strategy. In the second scenario (c, d), the benefit of mutualism outweighs the cost regardless of the opposing player’s strategy. In the third scenario (e, f), the benefit of mutualism outweighs the cost only when the opposing player is a non-mutualist. Results are shown specifically for *x*^*^ = 0.5 (*c*_*t*_ = 1 and *B*_*MN*_ = 4) but generally apply to 0 > *x*^*^ > 1. For the best response curves (a, c, e), *x*_*i*_ indicates the best strategy for the ii-th player with greater values of *x*_*i*_ indicating mutualism. Solid lines are the best response for player 1 and dashed lines for player 2. As this is an intraspecific evolutionary game of a single population, the dotted line *x*_1_ = *x*_2_ indicates the feasible set of solutions. Actual solutions for *x*^*^ are the intersection of all three lines. (a) The best response leads to a single strategy ESS of non-mutualism fixation. (c) The best response leads to a single strategy ESS of mutualism fixation. (e) The best response leads to a multiple strategy ESS of coexistence between mutualism and non-mutualist types. Replicator dynamics show the same results as the best response curves (b, d, f); the only difference is that fixation of either strategy is an equilibrium in all three scenarios but the stability of those two equilibria varies according to the cost-benefit ratio.

**Fig. 2.**
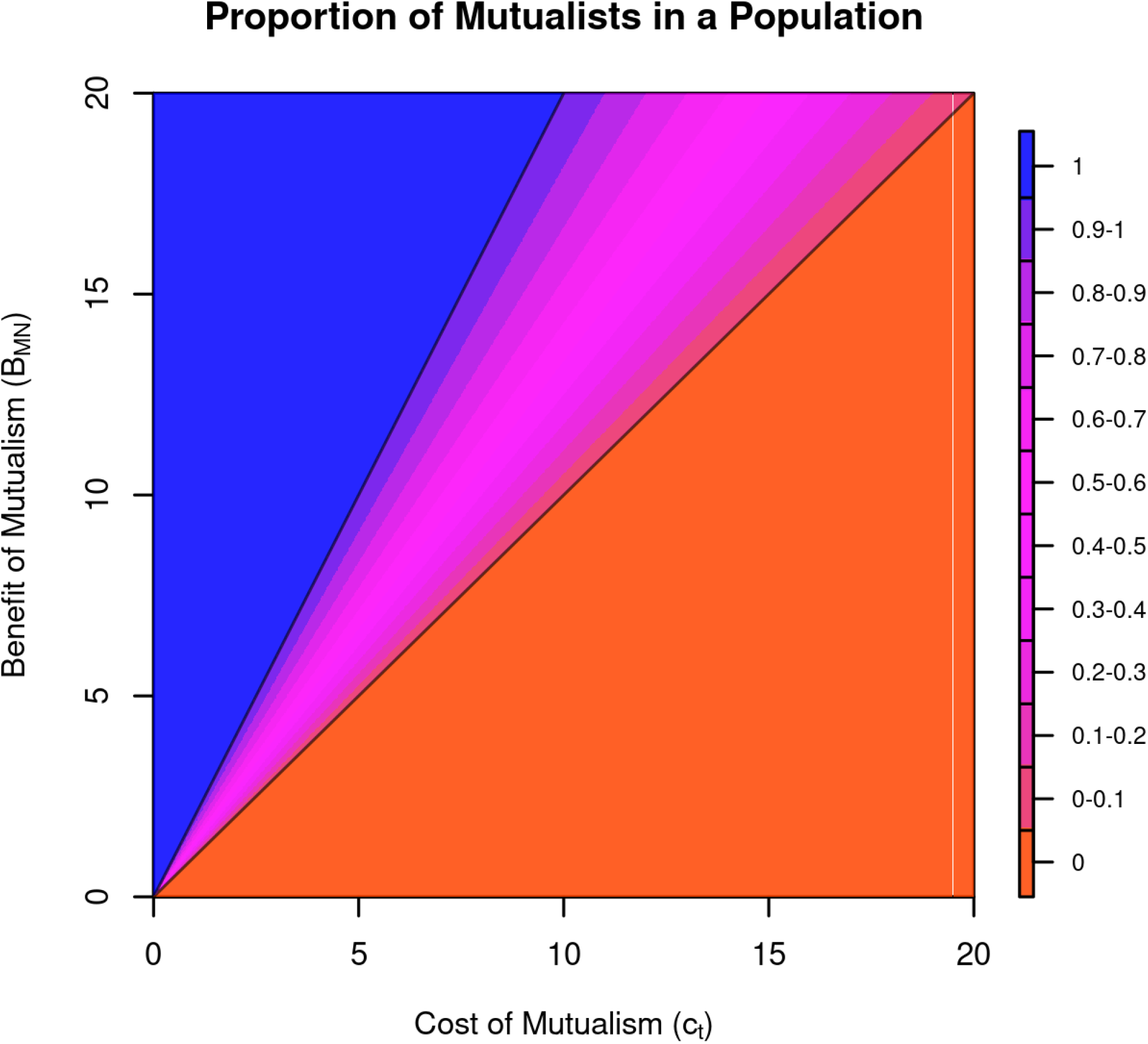
A plot of the proportion of mutualists in a population *x*^*^ for combinations of *B*_*MN*_ and *c*_*t*_. Orange-red indicates non-mutualist fixation, blue indicates mutualist fixation, and magenta indicates coexistence.

### Competition and neighborhood size

Above, we assumed that plants competed with only one other individual at a given time. While the non-motile nature of plants means that they compete on local spatial scales, this neighborhood of competitive interactions is generally more than one neighbor. It can be especially true when nutrients are scarce and multiple individuals must draw from the same pool leading to each individual taking up a smaller share of nutrients. For a mutualist plant, its share of the microbially available nutrients will also depend on the frequency of mutualists in the neighborhood which ultimately depends on the frequency of mutualists in the population. Therefore, we modify our game to have a plant compete with any number of individual plants in its local neighborhood. We can generalize our fitness matrix such that

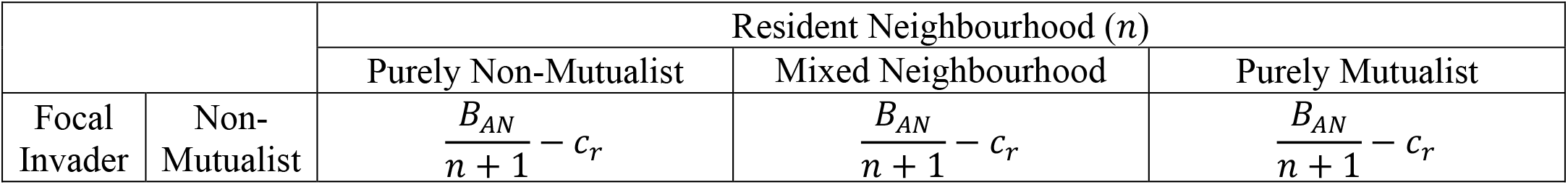
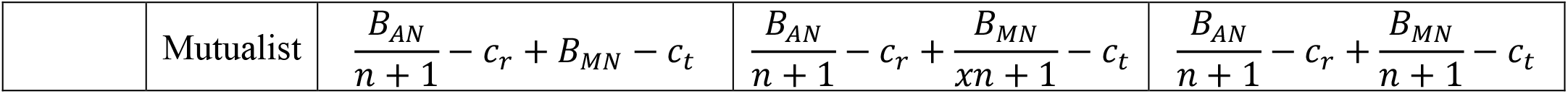

| | | Resident Neighbourhood ($n$) | | |
| --- | --- | --- | --- | --- |
|  |  | Purely Non-Mutualist | Mixed Neighbourhood | Purely Mutualist |
| Focal Invader | Non-Mutualist | $\frac{B_{AN}}{n+1} - c_r$ | $\frac{B_{AN}}{n+1} - c_r$ | $\frac{B_{AN}}{n+1} - c_r$ |
| | Mutualist | $\frac{B_{AN}}{n+1} - c_r + B_{MN} - c_t$ | $\frac{B_{AN}}{n+1} - c_r + \frac{B_{MN}}{xn+1} - c_t$ | $\frac{B_{AN}}{n+1} - c_r + \frac{B_{MN}}{n+1} - c_t$ |

where *n* is the number of competitors per plant, i.e., the size of its local neighborhood, and *x* is the proportion of mutualists in that neighborhood. Like before, fitness benefits from freely available nutrients are invariant with strategy. Therefore, it can be subtracted from each expression to arrive at the simpler matrix below.

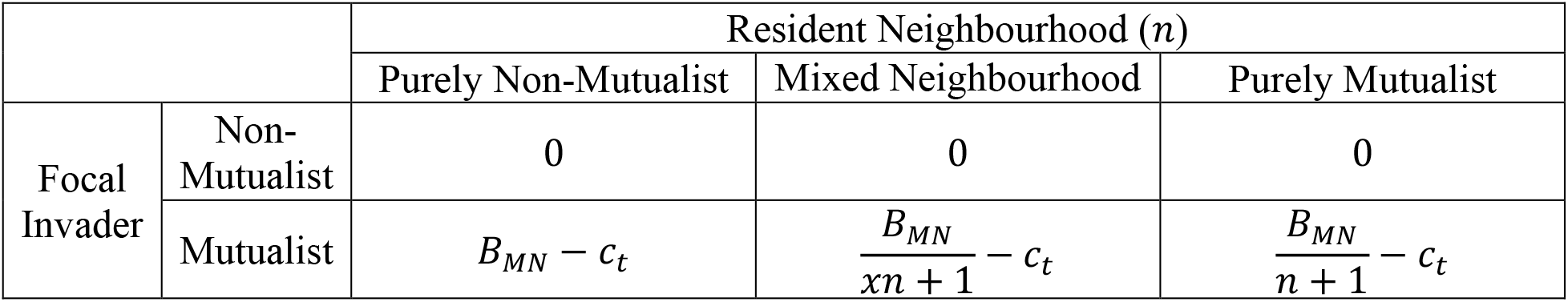

Following from Hauert et al.(2006), we derive overall fitness of a mutualist plant to be 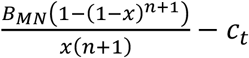 assuming local neighborhoods are generated randomly (see SI for derivation). With a larger neighborhood of interaction, the criterion for non-mutualist fixation is unchanged and still requires that the cost of mutualism without mutualist competitors must be greater than the benefits *c*_*t*_ > *B*_*MN*_. Fixation of the mutualist strategy requires that the benefit of mutualism when solely competing with mutualist must be greater than the costs 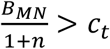. We can express this criterion in terms of a cost-benefit ratio 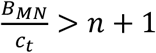. From this ratio, we can see that as *n* increases, there needs to be more benefits relative to the costs, reducing the possibility of fixation. This means that mutualist strategy is more likely to appear in coexistence with the non-mutualist strategy with an increasing number of competitors (Figure 3). Solving for this coexistence equilibrium proportion of mutualists is significantly harder with multiple competitors, and is analytically impossible with five or more individuals, but we can arrive at the solution 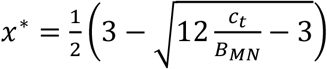 when there is a neighborhood of two plant competitors (see SI for the solution for three competitors).

**Fig. 3.**
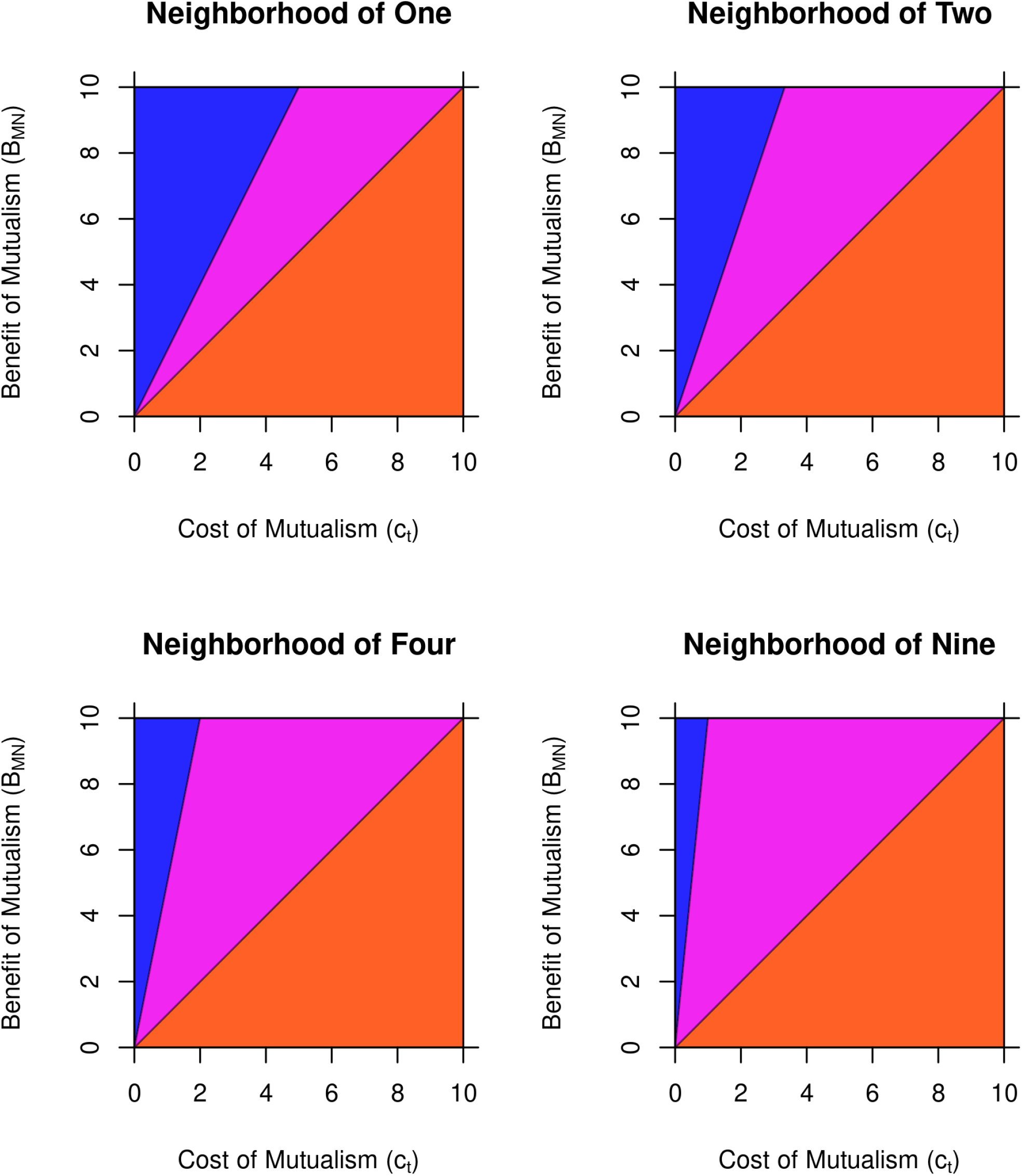
Plots of how regions of coexistence change with increasing neighborhood size. The colors remain the same as Figure 2. The region of fixation for the non-mutualist strategy does not change with neighborhood size and the same is true for the region where mutualist strategy is present, i.e., the combined region of mutualist fixation and coexistence. However, the region of mutualist fixation becomes smaller, expanding the region of coexistence between strategies.

## Discussion

In this study, we wanted to see how the ecological and environmental context in which a plant that partners with a microbe to obtain nutrients finds itself could lead to an intraspecific competitive advantage. Many models of mutualism evolution focus on the stability of the plant-microbe partnership, especially with regard to microbial cheating and the maintenance of beneficial variants (West *et al*. 2002; Akçay and Roughgarden 2007; Akçay and Simms 2011). Host-host interactions are usually not a focus in these models of evolution but rather are treated implicitly (Bergstrom and Lachmann 2003) (however see (Lu and Hedin 2019)). Our model explicitly focuses on host-host competition and the competitive advantage for a host plant. Using evolutionary game theory, we found the unsurprising result that if the cost of mutualism outweighed the benefit, then non-mutualists would entirely exclude mutualist while if the benefit of mutualism was greater than the cost under at least some conditions, then mutualism would be a viable strategy. That evolution favors traits with higher benefits compared to costs is well known, but by expanding the neighborhood size, we gained more precise insight into how benefits and costs combined within the context of intraspecific plant competition shape the evolution of mutualism. In particular, the evolution of mutualism was heavily influenced by the number of plants in the local neighborhood with which an individual would compete, as fixation of the mutualism strategy became harder with a larger local neighborhood, instead more often resulting in coexistence between mutualist and non-mutualist strategies. Alternatively, fixation of the non-mutualist strategy was invariant with the size of the neighborhood (Fig 3). Thus, our model predicts that mutualist and non-mutualists should frequently coexist within the same population and that the frequency of mutualists declines with the size of the local neighborhood.

Our model was simple. It assumed that the benefits and costs of obtaining nutrients were constant, only changing with competition between host plants. Because all host plants competed equally for the same freely available nutrients regardless of strategy, it had no effect on our results. All that mattered was the net benefit of mutualism. This conflicts with empirical studies that have shown that increasing nitrogen availability leads to a reduction in the mutualist partnership (Vitousek *et al*. 1997; Weese *et al*. 2015; McCoy *et al*. 2018; Taylor and Menge 2018) (but see (Simonsen *et al*. 2015)). This suggests that microbial mutualism does not simply occur as an added benefit to the plant. Instead, there must be some tradeoff between using freely available nutrients and microbially obtained nutrients. This could be due to a fixed resource budget on the part of the plant – anywhere between 4% and 20% of total plant carbon is traded to mychorrhizal partners (Johnson *et al*. 1997; Voisin *et al*. 2003; Taylor and Menge 2018) – varying marginal costs of investment in the sources of the nutrients, preference for the form the nutrient comes in (Falkengren-Grerup 1995), or some combination of the three.

One interesting result of our model is that coexistence only happened if mutualists competed for the same microbially obtained nutrients. If they did not compete, then it would lead to fixation of either strategy as either could be competitively dominant. We know that some microbial mutualisms differ in their nutrient sources. Mycorrhizae obtain their traded nutrients such as phosphorous and nitrogen from organic sources (Hawkins *et al*. 2000; Hodge *et al*. 2001; Leigh *et al*. 2009), a depletable resource likely shared between mutualist competitors. Rhizobia, on the other hand, get their traded nitrogen from fixing atmospheric nitrogen, a functionally unlimited resource that likely is not locally depletable (Sessitsch *et al*. 2002; Sawada *et al*. 2003). In the rhizobial mutualism, benefits may not change in the presence of competitors with the same strategy. This lack of sharing the microbially derived resources may add to the explanation as to why legumes are so dominant in mutualistic invasions compared to mycorrhizal associated plants (Richardson *et al*. 2000; Castro-Díez *et al*. 2014). If a mutualist invader must share its resources with other competitors, it becomes limited by its own success; with more individuals using the same strategy, frequency dependence puts an upper limit on how successful an invader can be, especially with a larger neighborhood of competition. By not having to share resources, invading legumes may represent a purely dominant strategy, at least in the right conditions.(de Faria *et al*. 1989; Simon *et al*. 1993; Sprent 2005; Wang and Qiu 2006; Werner *et al*. 2015)(Maherali *et al*. 2016; Lu and Hedin 2019)(Downie 2014; Hoffman *et al*. 2014)

Modifications to this model can be made to reveal other aspects of mutualism evolution. For example, we assumed that a plant either was a mutualist and so fully invested in mutualism or was not a mutualist regardless of whether net benefits were positive or negative. This is likely true at larger scales and interactions at the intertaxonomic level where entire lineages show the presence or absence of mutualism strategies (Sprent 2005; Werner *et al*. 2015). However, at smaller scales of the individual and population, variation in mutualism is likely to present itself in a more continuous and quantitative fashion (Heath and Stinchcombe 2014). The abstract nature of mathematical modelling does mean that our equilibrium proportion *x*^*^ could be understood as the proportion of mutualists in a population or community depending on whether the interactions are thought to be intra-or interspecific respectively as well as probability of any individual using the mutualism strategy. However, different processes and properties operate on these different scales (Jablonski 2008). At the individual level, timescales are within a lifetime, and the response is governed at the anatomy and physiology specific to that organism. At the population and community level, timescales operate over generations with variation between individuals leading to variations in fitness and reproduction which drive the response. Both scales are unique but influence each other; seeing how plasticity at the individual level drives variation at the population/community level and vice versa would certainly reveal much about the dynamics of mutualism evolution. Such a model of plasticity in the amount of trade would require more than just fitness benefits of nutrients, it would require a second resource (i.e. carbon) for the plant to trade. We suggest that this model could become a more process-based model of plant growth that includes photosynthesis to acquire carbon for trade as well as nutrient dynamics in soil. A number of models of plant growth with limitation from multiple essential resources exist (Pacala and Tilman 1994; Craine *et al*. 2005; Dybzinski *et al*. 2011; McNickle *et al*. 2016). Future work could explore introducing some of the insights gained in our simple model into those more complex models of plant growth and allocation.

This simplicity of our model does offer an advantage in that it can be easily translated to an experimental setup for falsification. One potential set up could be pot experiments with mutualist and non-mutualist varieties of plants (McNickle et al., 2020). Some plant species have loss of function mutants that allow for resource mutualisms to be turned on or off such as *DMI1* in *Medicago* and *sym8* in *Pisum* (Markwei and LaRue 1992; Balaji *et al*. 1994; Guinel and Geil 2002; Ané *et al*. 2004). One could grow the mutants and wildtype of the same species together in the same space with different densities and nutrient concentrations to see how they respond.

Fitness proxies like seed and flower number, average seed size, plant height, and root and shoot biomass could be measured for comparisons between wildtype and mutants (subsequent statistical analyses would have to take into account intrinsic fitness differences between wildtype and mutant as wildtype typically have greater fitness than mutants). Because these mutants do not express mutualisms with both mycorrhizae and rhizobia, comparisons between different microbial partners can also be made.

## Conclusion

Our model, though simple, reveals that a host can gain a competitive advantage from partnering with a microbe, leading to the evolution of mutualism in a population and fixation in a lineage. It also points to the possibility of coexistence of mutualist strategies in a population, an experimentally testable hypothesis. The results elucidate the basic conditions of positive net benefit and low local competition needed for this competitive advantage, why mutualisms may be prevalent yet variable, and how this prevalence and variation depends on sharing of resources. We suggest that future models incorporate mutualism into process-based models of plant growth.

## Acknowledgements

AHH’s contribution to this work was supported by the National Science Foundation Postdoctoral Research Fellowship in Biology Grant no. 2010972. GGM was supported by the USDA National Institute of Food and Agriculture Hatch project 1010722.

