## Supplementary Information for "When Does Mutualism Offer a Competitive Advantage? A Game-Theoretic Analysis of Host-Host Competition in Mutualism"

In this supplementary information, we show the derivation of analytical solutions for our evolutionary game as referenced in the main text.

*Competition with one plant*

From the main text, the payoff matrix for competition with one plant looks like this:

| | | Resident | |
|---|---|---|---|
| | | Non-Mutualist | Mutualist |
| Focal Invader | Non-Mutualist | 0 | 0 |
| | Mutualist | $B_{MN} - c_t$ | $\frac{B_{MN}}{2} - c_t$ |

We denote the equilibrium proportion of mutualists as $x^*$. In this world, the fitness of a mutualist individual is $(1 - x^*)(B_{MN} - c_t) + x^*\left(\frac{B_{MN}}{2} - c_t\right) = \frac{B_{MN}}{2}(2 - x^*) - c_t$. The fitness of a non-mutualist individual is 0. At equilibrium, the fitness of both individuals will be equal, and therefore simplifying the equation $\frac{B_{MN}}{2}(2 - x^*) - c_t = 0$ yields $x^* = 2\left(1 - \frac{c_t}{B_{MN}}\right)$.

*Competition and neighborhood size*

The payoff matrix for competition with multiple plants from the main text is:

| | | Resident Neighbourhood ($n$) | | |
|---|---|---|---|---|
| | | Purely Non-Mutualist | Mixed Neighbourhood | Purely Mutualist |
| Focal Invader | Non-Mutualist | 0 | 0 | 0 |
| | Mutualist | $B_{MN} - c_t$ | $\frac{B_{MN}}{xn + 1} - c_t$ | $\frac{B_{MN}}{n + 1} - c_t$ |

Following Hauert et al. (2006), the fitness of an individual is the probability that it finds itself in a neighborhood with a certain composition multiplied by the fitness of that individual given that neighborhood. Once again, let $x^*$ be the equilibrium proportion of mutualists in a population. The probability that an individual finds itself in a neighborhood of $n$ individuals with $i$ mutualist individuals is $\binom{n}{i}(1 - x^*)^{(n-i)}(x^*)^i$ while its fitness in this neighborhood is $\frac{B_{MN}}{i+1} - c_t$ leading to a probabilistic fitness of $\binom{n}{i}(1-x^*)^{(n-i)}(x^*)^i\frac{B_{MN}}{i+1}-c_t$. This probabilistic fitness is summed over all probabilities $\sum_{i=0}^{n}\binom{n}{i}(1-x^*)^{(n-i)}(x^*)^i\frac{B_{MN}}{i+1}-c_t$ yielding $\frac{B_{MN}\left(1-(1-x^*)^{n+1}\right)}{x^*(n+1)}-c_t$. The fitness of a non-mutualist individual remains 0.

Because the fitness term is a higher order polynomial, we cannot get an explicit solution for all neighborhood size $n$. However, smaller neighborhoods can yield explicit solutions. We already derived the solution with a neighborhood of one, $n=1$, above. For a neighborhood of two, the fitness of a mutualist is $\frac{B_{MN}}{3}\left(3-3x+{x^*}^2\right)-c_t$. Setting this equation equal to zero (the fitness of the non-mutualist), using the quadratic equation, and selecting only valid solutions give us an equilibrium proportion of mutualists of $x^*=\frac{1}{2}\left(3-\sqrt{12\frac{c_t}{B_{MN}}-3}\right)$. For a neighborhood of three, the fitness of the mutualist is $\frac{B_{MN}}{4}\left(4-6x^*+4{x^*}^2-{x^*}^3\right)-c_t$ and the equilibrium proportion in the population is $x^*=\frac{1}{3}\left(4+\sqrt[3]{\frac{10B_{MN}-54c_t+6\sqrt{3B_{MN}^2-30B_{MN}c_t+81c_t^2}}{B_{MN}}}-\sqrt[3]{\frac{4B_{MN}}{5B_{MN}-27c_t+3\sqrt{3B_{MN}^2-30B_{MN}c_t+81c_t^2}}}\right)$. A neighborhood of four can also be solved using the same method. However, this derivation is extremely long and not particularly informative. Thus, we do not show it here.

**Hauert C, Michor F, Nowak MA, Doebeli M**. **2006**. Synergy and discounting of cooperation in social dilemmas. *Journal of Theoretical Biology* **239**.
